# Evaluation of *in vitro* activity of copper gluconate against SARS-CoV-2 using confocal microscopy-based high content screening

**DOI:** 10.1101/2020.12.13.422548

**Authors:** Killian Rodriguez, Rigaill Josselin, Estelle Audoux, Florian Saunier, Elisabeth Botelho-Nevers, Amélie Prier, Yann Dickerscheit, Sylvie Pillet, Bruno Pozzetto, Thomas Bourlet, Paul O. Verhoeven

## Abstract

**Context:** Severe Acute Respiratory Syndrome Coronavirus 2 (SARS-CoV-2) that emerged late in 2019 is the etiologic agent of coronavirus disease 2019 (Covid-19). There is an urgent need to develop curative and preventive therapeutics to limit the current pandemic and to prevent the re-emergence of Covid-19. This study aimed to assess the *in vitro* activity of copper gluconate against SRAS-CoV-2.

**Methods:** Vero E6 cells were treated with copper gluconate 18 hours before infection. Cells were infected with a recombinant GFP expressing SARS-CoV-2. Infected cells were maintained in fresh medium containing copper gluconate for an additional 48-hour period. The infection level was measured by the confocal microscopy-based high content screening method. The cell viability in presence of copper gluconate was assessed by XTT assay.

**Results:** The viability of Vero E6 cells treated with copper gluconate up to 200 μM was found to be similar to that of untreated cells, but it dropped below 40% with 400 μM of this agent. The infection rate was 23.8%, 18.9%, 20.6%, 6.9%, 5.3%,5.2% in cells treated with 0, 2, 10, 25, 50 and 100 μM of copper gluconate respectively. As compared to untreated cells, the number of infected cells was reduced by 71%, 77%, and 78% with 25, 50, and 100 μM of copper gluconate respectively (p < 0.05).

**Conclusion:** Copper gluconate was found to mitigate SARS-CoV-2 infection in Vero E6 cells. Furthers studies are needed to determine whether copper homeostasis could play a role in SARS-CoV-2 infection.

**GRAPHICAL ABSTRACT:** 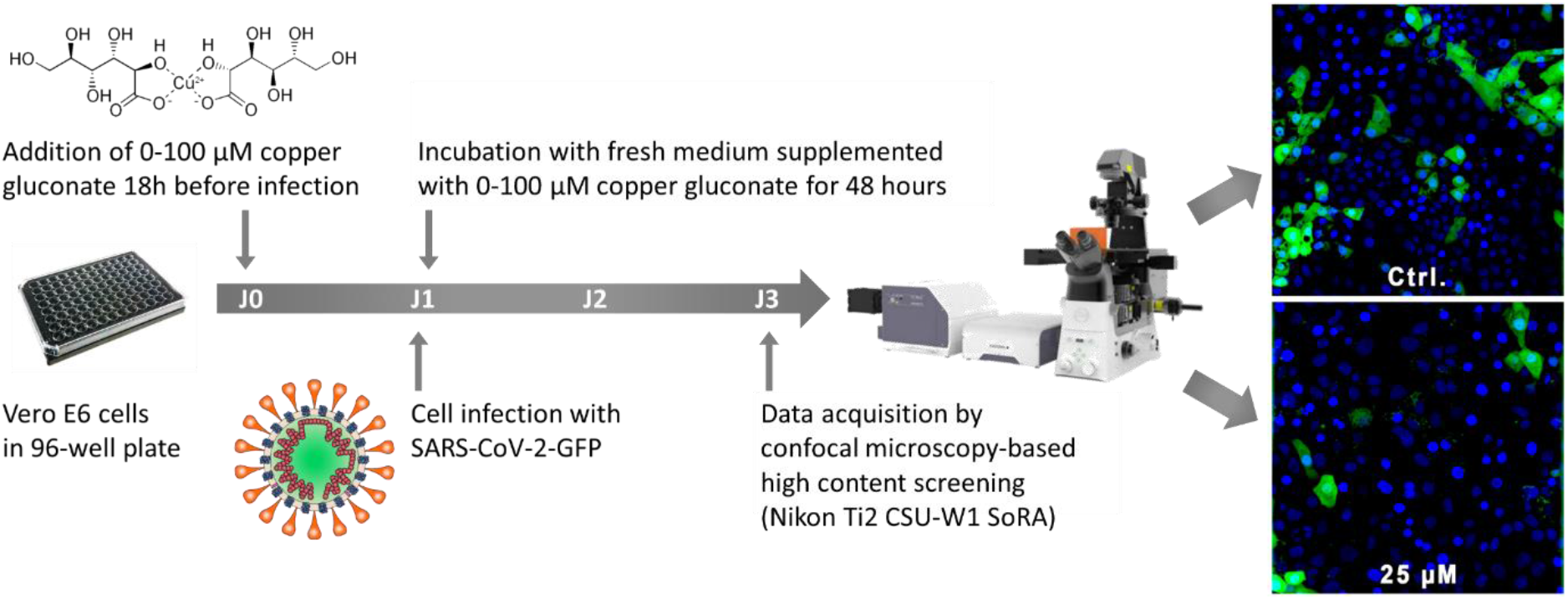

## INTRODUCTION

At the end of 2019, the emergence of a novel coronavirus designated as Severe Acute Respiratory Syndrome Coronavirus 2 (SARS-CoV-2) has led to a pandemic that threatens human health and public safety [1,2]. This new virus is highly transmissible and has spread very fast all over the world [1]. Even though the great majority of people (i.e. around 80%) develop mild to moderate coronavirus disease 2019 (Covid-19), a significant proportion of cases are severe or critical and can lead to death [1,3]. So far 1,374,985 people died from Covid-19 worldwide as of November 22th, 2020 [4]. Thus, there is an urgent need to contain SARS-CoV-2 spread and virulence with effective curative and preventive treatments [2,3].

During the early phase of the SARS-CoV-2 outbreak, we have been faced with a strong need for *in vitro* models able to test the efficacy of compounds against this virus but only a few laboratories were able to do so. *In silico* studies have identified dozens of drugs potentially active against SARS-CoV-2 [5–10]. Despite drug repurposing has been considered as one of the most promising strategies for improving the care of Covid-19 patients, published data on the *in vitro* efficacy of molecules potentially active on SARS-CoV-2 remain very limited to date [11–14]. Available studies focused mostly on few drugs including hydroxychloroquine, remdesivir, lopinavir, ritonavir, interferon, umifenovir, favipiravir, camostat mesylate, and immunomodulatory therapies [10–14].

Although some treatments have shown some benefits in patients at later stages of the disease (i.e. dexamethasone, anticoagulation treatments), there are no acknowledged effective antiviral therapies for Covid-19 [1]. Recently, preliminary results of the SOLIDARITY trial showed that hydroxychloroquine, remdesivir, lopinavir/ritonavir, and interferon regimens have no significant effect on the mortality rate nor on the length of hospital stay in COVID-19 patients [15]. Other ongoing clinical trials could provide additional results shortly [16].

Among the other compounds found to be directly active against SARS-CoV-2, essential minerals may require special attention. Antimicrobial and antiviral activity of copper is well established [17]. Copper ions have been found to elicit a broad action against viruses including coronaviruses [18–21]. Recently, it has been shown that SARS-CoV-2 can be eradicated from a copper surface within 4 hours while it can survive up to 72 hours on stainless steel and plastic surface [22]. Copper has been proposed to prevent transmission of the SARS-CoV-2 in the hospital environment (i.e. to cover door handles) or in application to face masks with the aim of reducing the risk of catching or spreading SARS-CoV-2 [21,23].

In eukaryotes, copper acts as an essential cofactor for more than 30 enzymes involved in redox reactions including superoxide dismutase (SOD) and ceruloplasmin. As well as for other trace metal ions (*e.g*. iron, manganese, zinc, selenium, and cobalt), maintenance of an adequate intracellular concentration of copper is essential to avoid the negative metabolic effects [24,25]. In humans, the normal cupremia varies from 9.75 to 27.75 μmol/L (650 to 1850 μg/L) in adults [24,26–28] and in tissues copper concentration fluctuates from 1 to 12 μg/g of tissues [24,28,29]. In human cells, copper is internalized by copper transporters 1 and 2 and is used for the synthesis of copper-requiring enzymes; it is stored mainly in the mitochondria and secreted by cells in the bloodstream; excess of copper is mostly eliminated by hepatocyte in bile [24]. In physiological conditions, copper is bound to ceruloplasmin (accounting for 40-70% of total plasma copper), albumin, alpha-2 macroglobulin, and other copper-carrying proteins for avoiding uncontrolled redox activity [24,29].

To the best of our knowledge, no published study to date have evaluated the effect of copper gluconate using an *in vitro* cell model of SARS-CoV-2 infection. This study aimed to assess if pre- and post-exposure treatment with copper gluconate could prevent the cells to be infected. For this purpose, we developed an original confocal microscopy-based high content screening (HCS) method using a recombinant GFP expressing SARS-CoV-2.

## METHODS

### Cells lines

Vero E6 and BHK-21 cells were cultured in Dulbecco’s modified Eagle’s medium (DMEM, ref. D6429, Sigma-Aldrich, Saint Quentin Fallavier, France) supplemented with 2% foetal bovine serum (FBS) (ref. 10270106, Gibco, ThermoFisher, Courtaboeuf, France) without antibiotic. All cells were maintained at 37°C and in a 5% CO_2_ atmosphere.

### Bacterial and yeast strains

*Escherichia coli* TransforMax Epi300 electrocompetent cells (Ref. EC300110, Epicentre, Madison, WI) and *Saccharomyces cerevisiae* VL6-48N strain [30] were used to propagate the pCC1BAC-His3 containing viral cDNA. *E. coli* bacteria were grown in LB broth supplemented with 25 μg/ml of chloramphenicol. *S. cerevisiae* yeast were grown on YPD agar supplemented with 25 μg/ml of chloramphenicol.

### Measurement of cell viability

To determine the toxicity of the chemical compound, cells were exposed to different concentrations of copper gluconate (Copper di-D-gluconate, CAS number 527-09-3, Isaltis, Lyon, France) ranging from 0 to 1600 μM for 24 hours in DMEM 2% FBS. Cell viability was determined in three independent experiments with the CyQUANT XTT assay (ref. X12223, Invitrogen, ThermoFisher) following the manufacturer’s recommendations. Optical densities at 450 and 660 nm were measured using a microplate reader (Sunrise, Tecan, Lyon, France).

### Rescue of synSARS-CoV-2-GFP

The BAC containing viral cDNA of synSARS-CoV-2-GFP clone 6.2 was kindly provide by Volker Thiel [31]. Upon receipt, BAC was stored in *S. cerevisiae* VL6-48N. The YAC DNA was extracted using spin column-based nucleic acid purification (ZR BAC DNA Miniprep Kit, ref. D4049, Zymo Research, Irvine, CA). The YAC containing viral cDNA was transformed into *E. coli* TransforMax Epi300 and amplified flowing manufacturer recommendations. The BAC was extracted from *E. coli* TransforMax Epi300 using spin column-based nucleic acid purification (ZR BAC DNA Miniprep Kit) and stored at 2-8°C until used in the next days. The BAC containing viral cDNA was cleaved at a unique restriction site located downstream of the 3’-end poly(A) tail using NotI-HF (Ref. R3189S, NEB, Grenoble, France). In parallel, the N gene was amplified from the BAC containing viral cDNA with a high fidelity polymerase (Taq Q5 Hot Start High-Fidelity 2x Master Mix, Ref. M0494S, NEB) and primers PV012-F (5’-ACT-GTA-ATA-CGA-CTC-ACT-ATA-GGG-ATG-TCT-GAT-AAT-GGA-CCC-CAA-AAT-C-3’) and PV013-R (5’-GGC-CGC-GGC-CGC-TTT-TTT-TTT-TTT-TTT-TTT-TTT-TTT-TTT-TAG-GCC-TGA-GTT-GAG-TCA-GC-3’). *In vitro* transcription of 1-2 μg of phenol-chloroform extracted and ethanol precipitated DNA resolved in nuclease-free water was carried out using T7 RiboMAX expression large scale RNA production system (ref. P1300, Promega, Charbonnières-les-Bains, France) with m7G(5')ppp(5')G RNA Cap Structure Analog (ref. S1404L, Promega) as recommended by the manufacturer. A similar protocol was used to produce a capped mRNA encoding the N protein. Approximately 10 μg of *in vitro* transcribed viral genomic RNA was electroporated together with 2 μg of the N gene transcript into BHK-21 cells. Briefly, 10^7^ BHK-21 cells were resuspended in 350 μL of ice-cold PBS, mixed with RNA, and transferred to 0.2 cm electroporation cuvette. Electroporation was carried out with a Genepulser apparatus (ref. 1652660, Biorad, Marnes-La-Coquette, France) with one pulse of 140V and 25 msec. BHK-21 electroporated cells were immediately co-cultivated with 70-80% confluent Vero E6 cells in wells of 9.5 cm^2^. After an incubation of cells for 5 days, the cell supernatant was collected and cellular residues were removed by centrifugation at 3000g for 20 min. Clarified supernatant (passage 0) was stored and used to produce virus stocks for further analysis. All work involving the culture, production, and storage of SARS-CoV-2-GFP was performed in a biosafety level 3 (BSL3) laboratory.

### Confocal microscopy-based high content screening

Copper gluconate was dissolved in sterile water and filter-sterilized with a 0.22 μm PVDF filter. The stock solution at 10 mM was stored at 2-8°C for a maximum of 10 days. Vero E6 cells were seeded in a 96-well plate (ref. CLS3904, Corning, Sigma Aldrich) at a density of 20,000 cells per well. Cells were incubated for 18 hours at 37°C and in 5% CO_2_. The next day, cells were treated with copper gluconate concentrations ranging from 0 to 100 μM for 18 hours before infection with GFP expressing SARS-CoV-2. The medium was removed and cells were infected with GFP expressing SARS-CoV-2 at a m.o.i. of 0.005 for 1 hour. After the adsorption step, the medium was removed and cells were incubated in fresh medium (DMEM supplemented with 2% of FBS) supplemented with copper gluconate concentrations ranging from 0 to 100 μM at 37°C and in 5% CO_2_ for another 48 hour-period. Nuclei of Vero E6 cells were stained with 4 μg/ml Hoechst 33342 (ref. H1399, Invitrogen) for 30 min. Ninety-six-well plates were sealed in the BSL3 laboratory to be imaged by confocal microscopy at 40-fold-magnification (Ti2 CSU-W1 SoRA, Nikon, France). Three independent experiments were performed and z-stack images were acquired in 12 random fields in duplicate wells (i.e. 6 fields per well) for each experimental condition. The whole process of acquisition was fully automated using in house pipeline developed with the JOBS module of the NIS software. Images were analysed using NIS general analysis 3 in-house pipeline (NIS software v5.30, Nikon) to count the total number of nuclei, the number of nuclei in the infected area, the volume of the infected area, and the mean fluorescence intensity (MFI) of infected cells.

### Statistics and software

SnapGene software v5.2 (GSL Biotech, San Diego, CA) was used to determine *in silico* the restriction profiles of BAC/YAC containing viral cDNA of synSARS-CoV-2-GFP clone 6.2. Statistical analysis and graphics were computed with GraphPad software v9 (Prism, San Diego, CA). The improvement of the images for publishing was performed with Fiji software (v1.53c).

## RESULTS

### Cell toxicity of copper gluconate

Vero E6 cells were treated with gluconate copper concentrations ranging from 0 to 1600 μM for 24 hours. Cell viability was determined by measuring the reduction of XTT converted to orange-coloured formazan product using the CyQUANT XTT assay. The viability of Vero E6 treated with copper gluconate up to 200 μM was similar to that of untreated cells. However, the cell viability dropped below 40% at 400 μM and reached a value close to zero for a concentration of 800 μM (**Figure 1**).

**Figure 1.**
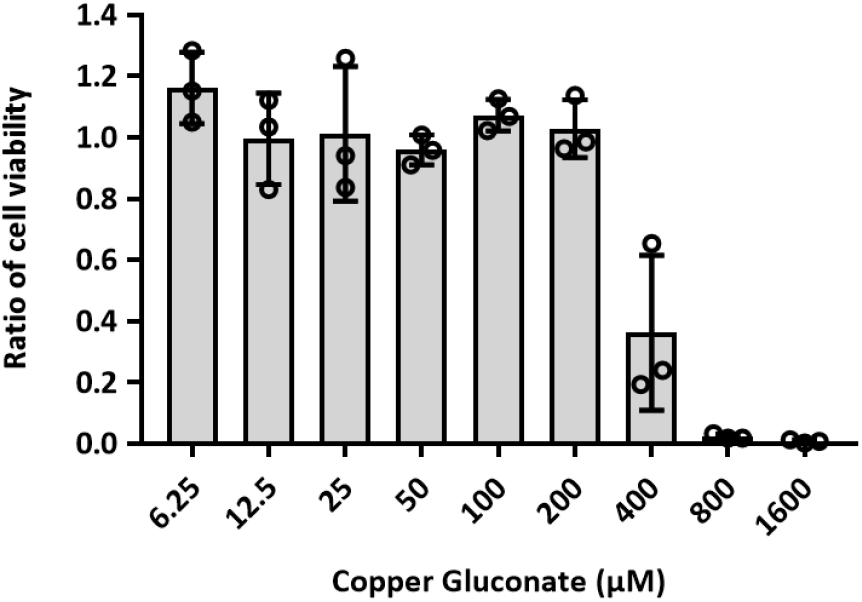
Viability of Vero E6 incubated with copper gluconate for 24 hours. The ratio of viable cells was calculated by dividing the signal of treated cells by the signal of untreated cells. Dots represent the value of each independent experiment, which correspond to the mean value of duplicate wells. Bars represent the mean value of three independent experiments. Error bars represent the standard deviation of the mean.

### Antiviral activity of copper gluconate

Vero E6 cells were treated with concentrations of copper gluconate ranging from 0 to 100 μM at 18 hours before SARS-CoV-2-GFP infection. The treatment was removed during the 1 hour of the adsorption of recombinant SARS-CoV-2-GFP viruses. After absorption, the medium was replaced by fresh culture media supplemented with the same concentrations of copper gluconate and cells were incubated for another 48 hour-period to let viruses infect and replicate in cells. Then, the level of infection was measured by confocal microscopy. Results of three independent experiments are depicted in **Figure 2**. The rate of infection was significantly lower in cells treated with a concentration of copper gluconate of 25 μM and higher as compared to untreated cells (**Figure 2a**). The rate of infection was 23.8%, 18.9%, 20.6%, 6.9%, 5.3%,5.2% in cells supplemented with 0, 2, 10, 25, 50 and 100 μM of copper gluconate respectively. Thus, the number of infected cells was reduced by 71, 77, and 78% with 25, 50, and 100 μM of copper gluconate respectively (p < 0.05). The cumulative number of cells analysed in three independent experiments represents roughly 5000 cells per condition tested in each independent experiment (**Figure 2b**). The number of infected cells was significantly lower in cells that were pre-treated with a concentration of copper gluconate of 25 μM and higher as compared to untreated cells (**Figure 2b**). The volume filled by infected cells was determined by measuring the volume filled by GFP-positive voxels for each independent experiment. Again, the concentration of copper gluconate of 25 μM was the lowest dose tested that significantly reduced the viral infection as compared to untreated cells (**Figure 2c**). Together, these results suggest that copper gluconate mitigate the infection of Vero E6 cells. Additionally, linear regression analysis showed that the MFI of infected cells decreased with the concentration of copper gluconate used to treat the cells (p < 0.01, linear regression) (**Figure 2d**). These latter results suggest that copper gluconate might also limit the viral replication inside Vero E6 cells.

**Figure 2.**
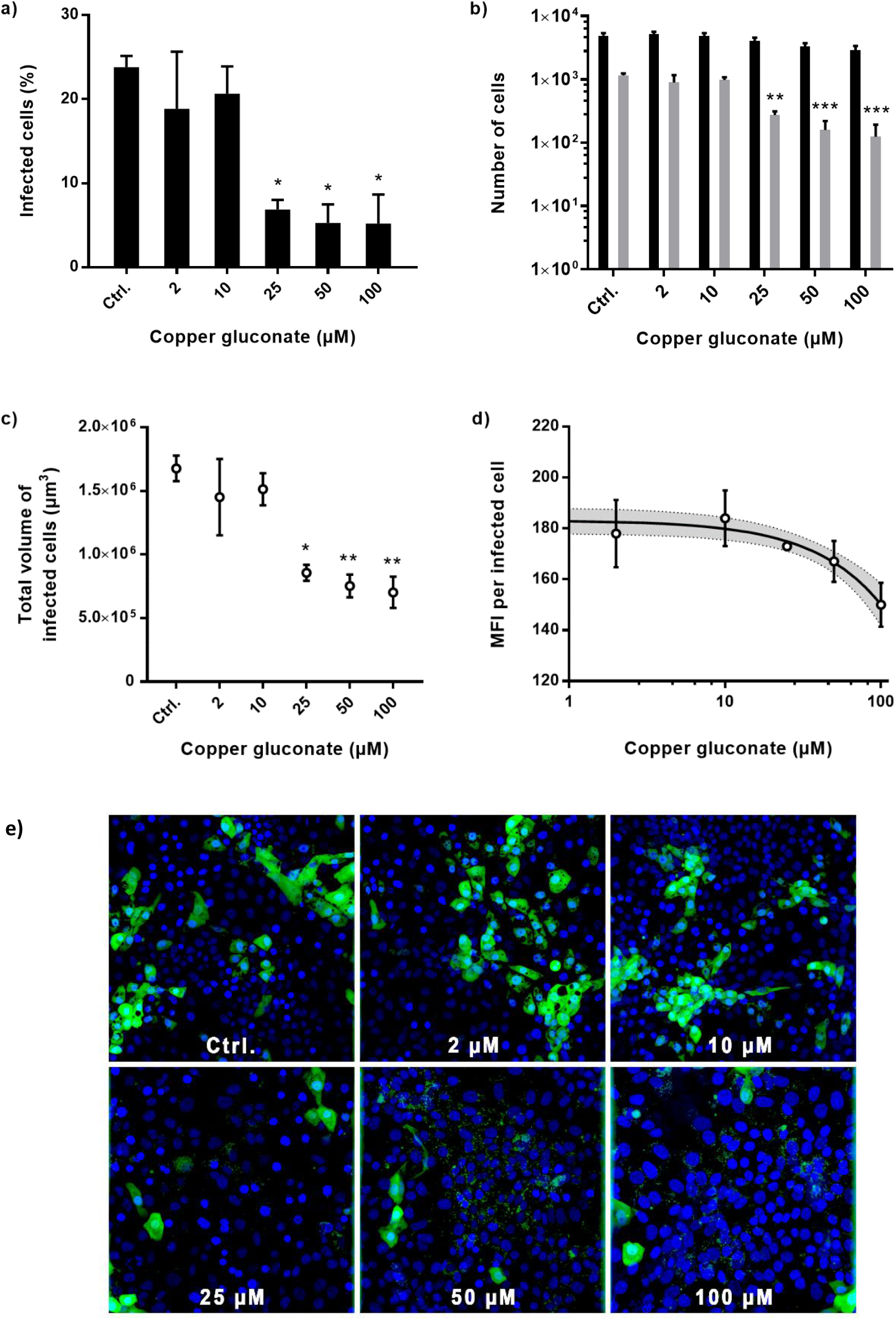
Antiviral effect of copper gluconate on Vero E6 cells infected with GFP expressing SARS-CoV-2 at 48 hours post-infection. Copper gluconate was incubated with Vero E6 cells for 18 hours before SARS-CoV-2 infection and until 48 hours post-infection. The confocal microscopy-based high content screening was used to measure the level of infection. Values represent the mean of three independent experiments. Error bars represent the standard error of the mean. **a)** Rate of SARS-CoV-2 infected cells. **b)** Number of cells (black bars) and cumulated SARS-CoV-2 infected cells (grey bars). **c)** Total volume filled by SARS-CoV-2 infected cells. **d)** Mean fluorescence intensity (MFI) per infected cell. Values represent the mean of three independent experiments. Solid line represents the linear regression (p < 0.01) with it 95% confidence interval (dots lines). **e)** Vero E6 cells (nuclei in blue) infected with GFP expressing SARS-CoV-2 (GFP in green) with copper gluconate ranging from 0 (Ctrl.) to 100 μM (40-fold-magnification). Images represent the projection of image stacks using the extended deep focus (EDF) algorithm with the NIS software (v5.30). EDF images were merged, false-coloured, and contrast-enhanced with ImageJ software (v1.53c) for display purposes. * p < 0.05; ** p < 0.01; *** p < 0.001; values were compared by one-way ANOVA with Dunnet correction for multiple comparisons for figures a), b) and c).

## DISCUSSION

For this study, we developed an original confocal microscopy-based HCS using an *in vitro* model with Vero E6 cells challenged with a recombinant GFP expressing SARS-CoV-2 that was reconstructed using a yeast-based reverse genetics platform [31]. This method has the advantage of being able to analyse each cell individually and to count thousands of cells per well at the same time to increase the reliability of the observations. By using both a motorized stage and a fully automated pipeline for image recording, we virtually eliminated any possible bias that could be linked to the person in charge of image acquisition. The quantitative analysis of images was also fully automated by using an in-house pipeline developed with a plugin of the NIS software to avoid technical bias. A similar experimental setup with Vero E6 cells and the same GFP expressing SARS-CoV-2 clone was found to be suitable for drug screening applications by using the antiviral remdesivir as a reference [31]. Thus, we decided to combine this *in vitro* model of GFP expressing SARS-CoV-2 infection with confocal microscopy-based HCS to assess the antiviral activity of copper gluconate. Furthers improvements of this technology (e.g. using cell lines with fluorescents reporters) could help to study more easily whether and how drugs or chemical compounds could counteract SARS-CoV-2 infection in mammalian cells.

In the present study, the combination of pre- and post-exposure treatment of Vero E6 cells with 25 to 100 μM of copper gluconate led to a 70% reduction of the cell infection rate. Copper gluconate concentrations of 25 μM and higher were also effective in reducing the number of infected cells and the volume filled by infected cells as compared to the untreated condition. However, none of the tested copper gluconate concentrations, up to 100 μM, was able to completely inhibit the infection. Although concentrations up to 200 μM showed no significant toxicity to Vero E6 cells, we did not use concentrations higher than 100 μM. In humans, the concentration of copper in whole blood is approximately 15 μM (1000 μg/L) but this value can fluctuate widely according to various factors [24,26–29]. In tissue, the copper concentration is about 1 to 12 μg/g, which is 1000-fold lower than the serum concentration [28,29]. It must be acknowledged that the effect we observed with 25 μM of copper gluconate is not as powerful as that described with antiviral drugs [32] but this concentration is reasonably close to the normal cupremia. The fact that a physiological concentration of copper was not able to abolish the viral infection is not so surprising if we consider the fact that copper is a ubiquitous trace element present in all eukaryotes. Even if copper seems to act against the virus by decreasing the number of infected cells, the antiviral effect observed is probably multifactorial in Vero E6 cells and certainly much more complex *in vivo*.

Warnes *et al.* showed that copper ions can damage virus membranes and destroy the viral genome of human coronavirus 229E [33]. In our study, a direct effect of copper cannot be excluded because copper gluconate was maintained in the culture medium as long as the infection lasted.

We also observed that the increase of copper gluconate concentration up to 100 μM is associated with a statistically significant decrease of MFI. Because GFP expressed by the recombinant SARS-CoV-2 is fused to the non-structural protein 7, we can speculate that copper might limit the synthesis of viral proteins. This hypothesis is supported by *in silico* studies predicting that metal ions such as cobalt(III) or copper(II) could inhibit the SARS-CoV-2 main protease [34,35]. However, found only two *in vitro* studies corroborating that copper ions could inhibit the synthesis of viral proteins or the replication cycle [36,37]. Thus, further *in vitro* experiments are needed to understand whether copper ions may limit the synthesis of viral proteins in mammalian cells.

Copper could also act against viruses by upregulating the Cu/Zn SOD1 expression. *In vitro* studies showed that an increase of SOD1 expression is associated with a decrease in viral replication [38,39]. Last, it was well established that the coronavirus replication complex requires autophagy-associated cellular components [40]. Because copper is known to be able to modulate autophagy, copper induced-autophagy could limit the availability of autophagy associated cellular components that are required for viral replication [19].

Whether one of these mechanisms more than another could be involved in the antiviral effect observed in our study remains unclear. Most proteins involved in copper homeostasis have not been studied extensively including ceruloplasmin and albumin, which have been the most researched [29]. Cuproenzymes such as ceruloplasmin or superoxide dismutase incorporate copper via the secretory pathway and they can be found in practically every location inside and out of the cell [41]. Further studies are needed to investigate whether the copper gluconate supplementation in culture media increases the internalization of copper and whether intracellular copper is required to struggle against viral infection.

In humans, a retrospective observational study showed that zinc and selenium transporter selenoprotein P and zinc deficiency was associated with the worst outcomes in elderly Covid-19 patients [42]. A meta-analysis in Chinese children reported that copper deficiency is associated with recurrent respiratory tract infection [43]. However, the knowledge about the pharmacokinetics of metal ions during the acute phase of viral infection is still limited [25] and whether copper homeostasis could play a role during SARS-CoV-2 infection is unknown. Thus, it could be interesting to determine copper concentrations in serum but also tissues such as nails and hairs in asymptomatic, mild, and severe Covid-19 patients. The animal models of SARS-CoV-2 infection that have been developed worldwide [44] seems the most appropriate approach to unravel the role of copper homeostasis during SARS-CoV-2 infection *in vivo*.

In conclusion, our findings showed that copper gluconate supplementation could reduce SARS-CoV-2 infection *in vitro*. Furthers studies are needed to determine whether copper homeostasis could play a role in SARS-CoV-2 infection.

## ACKNOWLEDGMENTS

The BAC/YAC containing the viral cDNA of the recombinant GFP-expressing SARS-CoV2 was kindly provided by Prof. Volker Thiel (Bern University, Switzerland). The authors acknowledge Nadine Ebert and Fabien Labroussaa (Bern University, Switzerland) for their helpful advice for the rescue of the recombinant GFP-expressing SARS-CoV-2. The strain of *Saccharomyces cerevisiae* VL6-48N was generously provided by Carole Lartigue-Prat (Bordeaux University, France). The Vero E6 cell line was a gift of Prof. Bernard La Scola (Aix-Marseille University, France). Christophe Machu and Ibrahim Hassani from Nikon Company are acknowledged for their technical assistance and for the helpful discussion regarding the analysis of image stacks. Copper gluconate used for this study was provided free of charge by EA Pharma company.

## FUNDING

This research was funded by the University Jean Monnet of St-Etienne (emergency financing for a microscope), the University Hospital of St-Etienne (donation from the St-Etienne football club), and EA Pharma company. This latter company had no role in the study.

Conflict of interest: none to declare.

## ROLES IN THE STUDY

TB and POV designed the study. KR performed the rescue of recombinant SARS-CoV-2-GFP. JR, EC, YD, AP performed experiments with infected cells. AP performed cytotoxicity assays. POV and JR developed the pipeline and analysed the data. FS, TB, and POV wrote the manuscript. All authors reviewed the manuscript.

